# A tension-induced morphological transition shapes the avian extra-embryonic territory

**DOI:** 10.1101/2024.02.08.579502

**Authors:** Arthur Michaut, Alexander Chamolly, Aurélien Villedieu, Francis Corson, Jérôme Gros

**Affiliations:** Institut Pasteur, Université de Paris, CNRS UMR3738, Developmental and Stem Cell Biology Department, F-75015 Paris, France; Laboratoire de Physique de l’Ecole Normale Supérieure, CNRS, ENS, Université PSL, Sorbonne Université, Université de Paris, 75005 Paris, France

## Abstract

The segregation of the extra-embryonic lineage is one of the earliest events and a key step in amniote development. Whereas the regulation of extra-embryonic cell fate specification has been extensively studied, little is known about the morphogenetic events underlying the formation of this lineage. Here, taking advantage of the amenability of avian embryos to live and quantitative imaging, we investigate the cell- and tissue-scale dynamics of epiboly, the process during which the epiblast expands to engulf the entire yolk. We show that tension arising from the outward migration of the epiblast border on the vitelline membrane stretches extra-embryonic cells, which reversibly transition from a columnar to squamous morphology. The propagation of this tension is strongly attenuated in the embryonic territory, which concomitantly undergoes fluid-like motion, culminating in the formation of the primitive streak. We formulate a simple viscoelastic model in which the tissue responds elastically to isotropic stress but flows in response to shear stress, and show that it recapitulates the flows and deformation of both embryonic and extra-embryonic tissues. Together, our results clarify the mechanical basis of early avian embryogenesis and provide a framework unifying the divergent mechanical behaviors observed in the contiguous embryonic and extra-embryonic territories that make up the epiblast.

**Highlights:**

- The extra-embryonic region expands radially during epiboly
- Cell area increase accounts for the rapid extra-embryonic expansion
- Epiboly-induced tension reversibly stretches extra-embryonic cells
- A simple viscoelastic model recovers the morphogenesis of the entire epiblast

## Introduction

The evolution of extra-embryonic tissues supporting vertebrate development independently of the aquatic environment is a major innovation that enabled the elaboration of the cleidoic egg and the successful terrestrial colonization of amniotes. In avians, extra-embryonic tissues can be traced back to egg deposition. At this stage, the blastoderm consists of the *area pellucida*, a central transparent epithelial disk whose cells give rise to the embryo proper (EP), surrounded by the *area opaca*, a contiguous ring of optically opaque cells, that will give rise to the extra-embryonic (EE) yolk sac. Upon incubation, gastrulation begins; the contraction of supracellular actomyosin cables developing at the margin between the EP and EE territories transforms an initially crescent-shaped region into the primitive streak. A fluid-like response of the EP enables the tensile forces of the margin to shape the entire embryonic region (*1*). Concomitantly, the blastoderm expands until it eventually engulfs the entire yolk (Fig. 1A). This process named epiboly is conserved between fish, amphibians, reptiles, birds, and egg-laying mammals. Studies in fish have shown that epiboly is driven by a contractile actomyosin belt in the yolk syncytial layer (YSL) (*2*). A combination of circumferential contraction and friction force at this ring pulls the edge of the blastoderm over the entire YSL. Interestingly, although epiboly in fish and amphibians is tightly linked in space and time to the process of gastrulation, in birds and reptiles it operates independently, probably because of the larger yolk size that characterizes the cleidoic egg (*3*). Whereas the zebrafish blastoderm spreads over a 0.7mm diameter yolk within 5-6 hours, the chicken blastoderm engulfs a 5cm-diameter yolk over 4 days, leading to an over 1000-fold increase in blastoderm area. Early studies have shown that the processes driving avian epiboly differ from those identified in zebrafish and rely on the attachment of the rim of the blastoderm onto the inner surface of the vitelline membrane enveloping the yolk (*4*). These edge cells, which are the only ones in firm contact with the vitelline membrane, exhibit a specific molecular signature (*5*), and a distinct cellular organization (*6*) that allows them to adhere and migrate on the inner surface of the vitelline membrane. This migration has been described to generate tension across the entire blastoderm (*4, 7*). Several models have been proposed to explain how this tension drives the expansion of the blastoderm. It was first proposed that increased proliferation in the EE territory accounts for blastoderm expansion, and that the epiboly-induced tension only serves to arrange the epiblast, which would otherwise thicken into several layers, into a flat epithelial sheet (*4*). It was later suggested (based on measurements of the mitotic index across the blastoderm) that proliferation is uniform, and that the EE territory expands through cell flattening that is in part intrinsic to the cells and in part induced by tension (*8*). However, these hypotheses have never been tested and the contributions of proliferation, cell arrangements and stretching to the spreading of the epiblast, as well as their relation to the outward migration of its border, remain to be characterized. Importantly, when tension in the *area pellucida* is released by removing the *area opaca*, gastrulation is impeded (*7*). Altogether, these studies have suggested that the tension arising from the migration of the blastoderm edge is critical for the spreading of the EE territory and primitive streak formation in the embryonic region, yet the underlying mechanisms remain unknown.

**Figure 1.**
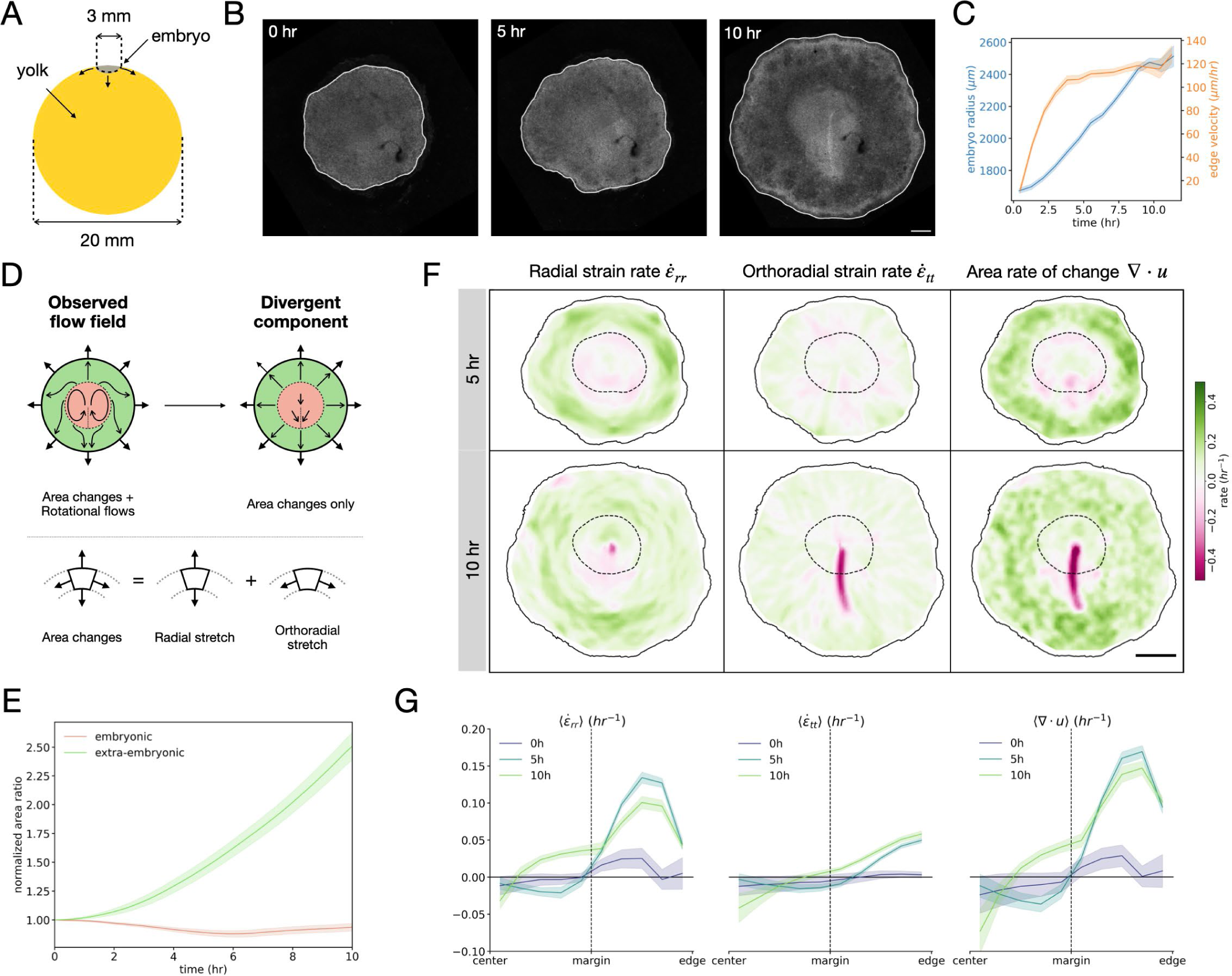
Epiboly is characterized by radial expansion of extra-embryonic territory. (A) Sketch of a quail epiblast on its yolk at the onset of epiboly (arrows). (B) Time series of a stage XI (EGK) transgenic memGFP embryo live-imaged for 10 hours. The white line shows the automatic segmentation of the epiblast’s edge. (C) Embryo radius and edge velocity measured by automatic segmentation averaged over N=11 embryos. (D) Upper panel: sketch of the extraction of the divergent component (area changes) from the tissue-velocity field. Lower panel: field decomposition into radial and orthoradial components. (E) Normalized evolution of embryonic and extra-embryonic areas, averaged over N=12 embryos. (F) Instantaneous maps of radial and orthoradial strain rate and total log area rate of change at 5 hours (top row) and 10 hours (bottom row) of embryo shown in (A). (G) Radial profile of the strain rate components and the total log area rate of change at 0, 5, and 10 hours, averaged over N=12 embryos. Shaded regions and error bars correspond to the standard error of the mean. Scale bars: 500 µm (B), 1 mm (F).

Although avian epiboly is an appealing *in vivo* model for the study of epithelial spreading, it has received surprisingly little attention, and has been characterized mostly using static, cellular, and molecular approaches. Here, combining dynamic imaging, transgenic quail embryos, quantitative analyses and mechanical approaches, we investigate the role of the epiboly-induced tension in the morphogenesis of the extra-embryonic and embryonic territories.

## Results

### Tissue-scale characterization of epiboly dynamics and associated extra-embryonic expansion

Epiboly has been previously characterized by manually measuring the entire epiblast disk area, at a few time points. To capture the dynamics of this process, we used the well-established *ex ovo* EC culture (*9*) to live-image stage XI (EGK) transgenic membrane-GFP (memGFP) quail embryos, every 6 min, over ∼12 hours, and quantified the velocity of the epiblast border by automatically segmenting the epiblast contour (Fig. 1B). Within 4 hours, the velocity of the epiblast edge gradually increased to reach an approximately constant plateau of 115 +/- 4 µm/hr (Fig. 1C, N=11 embryos). To verify that the *ex ovo* culture system does not interfere with epiboly, we also imaged an embryo directly in the egg. The dynamics of epiboly *in ovo* and *ex ovo* were very similar, with the epiblast edge reaching an approximately constant velocity of 144 µm/hr after a few hours (Fig. S1). Next, we characterized how the different embryonic territories deform as epiboly takes place. Embryonic and extra-embryonic territories have historically been defined using their optical density to transmitted light as the *area pellucida* and the *area opaca*, respectively. Although convenient, this method is imprecise and does not allow to identify and track these territories at later stages. As an alternative, we have previously shown that these territories can be recognized from the motion of the tissue, tracked using particle image velocimetry (PIV) (*1*). In addition, a mathematical decomposition of tissue flows into divergent (area changes) and rotational (incompressible shear) components was found to capture the differential deformation of the EP and EE territories. While this study showed that the incompressible rotational flows associated with primitive streak formation in the embryonic region can be quantitatively explained by a fluid-like yielding to tensile forces generated at its margin, it did not describe in detail the deformation of the EE territory nor address its mechanical control (for modeling purposes, the divergent component of the flow, treated as an intrinsic property of the different tissue regions, was taken from experiment). Here, we focus on the divergent flows (i.e. area changes) of the embryonic and extra-embryonic regions (Fig. 1D) and confirm that epiblast expansion is mostly due to area increase of the EE (Fig. 1E). To further quantify how tissues deform, we decomposed the local area expansion rate into radial 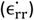 and ortho-radial 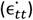 components (Fig. 1D, F). Whereas the EP maintains an approximately constant area over 12h, the area of the EE region increases as the tissue edge migrates outward. This growth is predominantly due to radial expansion, which like the edge velocity reaches a plateau at ∼5 hours, with ortho-radial expansion making a more limited contribution (Fig. 1F, Movie S1). Spatial mapping (Fig. 1G) revealed that the radial strain rate shows a radially symmetric annular pattern, peaking midway across the EE territory. The ortho-radial strain rate also exhibited a radially symmetric pattern, except at the primitive streak after the onset of area loss through cell ingression (Fig. 1G). Averaging strain rates over 12 embryos (based on common orthoradial landmarks, see methods) confirms the robustness of the annular pattern of radial expansion and the contribution of orthoradial expansion that peaks at the tissue edge (Fig. 1H). Altogether, these analyses identify a spatial pattern of radial expansion across the EE territory. Notably, these quantifications confirm that EP does not deform in the same way as EE, hinting that it is shielded from the influence of epiboly.

### Cellular basis of extra-embryonic expansion

We next asked how cells accommodate the rapid expansion of the EE territory. To this end, we performed high-resolution live-imaging of embryos carrying a memGFP reporter, which labels the cell-cell boundaries in the epithelial epiblast (Fig. 2A, B). To correlate the tissue and cellular scales, we followed regions of interest in these movies, computing their deformation over time using PIV (Fig. 2B, Movie S2) and automatically segmenting cell contours using a ‘Cellpose’ model (*10*) specifically trained on our dataset. This analysis showed that cell area expansion is a major contributor to tissue area expansion, and follows a similar pattern in space and time, peaking in regions midway across the EE (Fig. 2B, C-F, Movie S3), as seen in our analysis of tissue flows (cf. Fig. 1). Over time, the area of cells composing these regions increased by about a factor of 2.5-3, eventually reaching a plateau while tissue expansion proceeded (Fig. 2G). Quantifying several regions over multiple embryos (N=3 embryos) we found that over the course of 8 hours tissue area increases by roughly 4 fold, whereas cell areas increase on average by 2 fold (Fig. 2 H).

**Figure 2.**
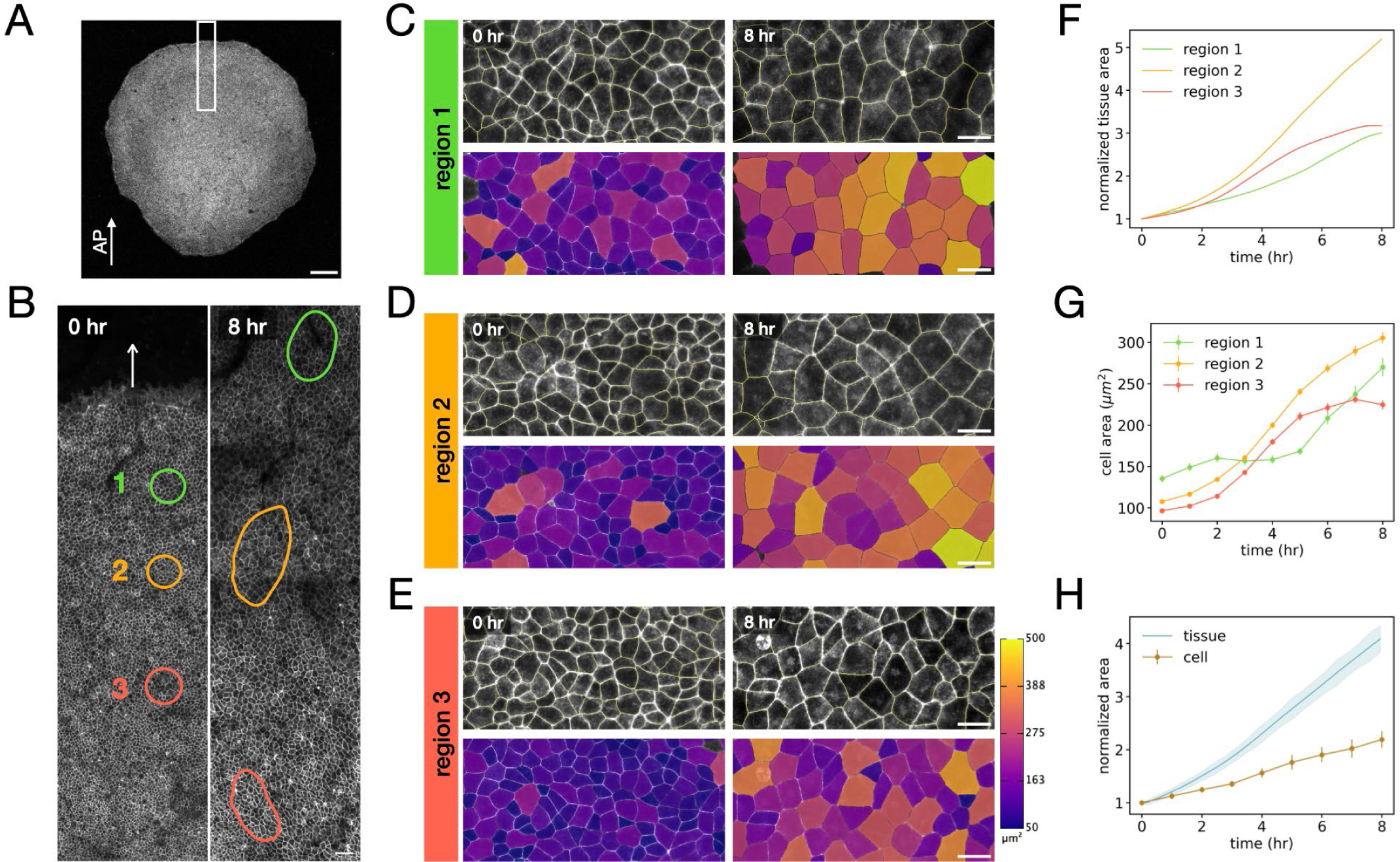
Cell area increase underlies extra-embryonic expansion. (A-B) High-resolution imaging of a strip of extra-embryonic tissue (white rectangle in [A]) in a memGFP embryo (AP, antero-posterior axis). Colored regions of interest in (B) are followed by PIV. The white arrow indicates the outward migration. (C-E) Time series of the PIV-registered regions of interest defined in (B). Top panels show the original image overlayed with the segmented cell contours as yellow lines. Bottom panels show the corresponding color-coded cell area quantification maps. (F-G) Evolution of region area (F) and average cell area within these regions (G) n=1691 cells, as defined in (B). (H) Evolution of region and cell area normalized and averaged over 8 regions from 3 embryos (N=4185 cells). Shaded regions and error bars correspond to the standard error of the mean. Scale bars: 500 µm (A), 50 µm (B), 20 µm (C-E).

Since cell area expansion on its own does not fully account for EE expansion, we considered the role of cell division. To this end, we first manually tracked a few cells and their progeny over time (Fig. 3A, Movie S4). Although cell areas overall increased at a steady pace, cell divisions led to an abrupt decrease in area (Fig. 3B). Still, daughter cells retained a larger area than that of their mother before epiboly and subsequently increased in area at a similar rate as before division (Fig. 2C, Fig. S2). These results suggest that EE expansion can be understood as the product of continuous cell area increase in an otherwise proliferating tissue. Of note, in these movies, few cell rearrangements could be observed over 3 hours, compared to the extensive rearrangements observed in the embryonic region over 1.5h (*11*). Although memGFP transgenic embryos reveal cell shape dynamics, automatic cell segmentation using membrane labeling is prone to errors that hinder the tracking of cells and their divisions over extended periods of time. Instead, to quantify more systematically the effect of cell division in EE tissue expansion, we live-imaged transgenic embryos expressing an H2B-GFP reporter gene. Nuclei labeled in this way can be very accurately segmented and tracked over time and the analysis of cell velocities and areas confirmed our PIV-based tissue-level tracking (Fig. S3). Combining PIV tracking of regions of interest with automatic segmentation of nuclei within these regions (Fig. 3D), we found that, as the regions are growing, cell numbers increase with a doubling time of 6.8 +/- 0.9 hr (Fig. 3 E). This estimate of the cell doubling time was confirmed by tracking individual cells that divided twice in the movie, yielding a cell cycle length of about 5.7 +/- 0.7 hr (Fig. 3E), in line with previous reports of cell cycle length in the chick epiblast (*12*). Finally, multiplying the average relative cell area change (from memGFP segmented cells) by the average relative increase in cell numbers (from segmented nuclei) recovered the time course of tissue expansion (from tissue-level tracking), confirming that our quantifications capture the contributions of cell division and area changes to extra-embryonic expansion (Fig. 3F). Altogether these analyses which quantify the effect of cell division confirm that cell area increase is an essential contributor to the rapid expansion of the EE territory. Whereas division does supply the tissue with new cells, the cell cycle ∼6h is too slow to accommodate the rapid areal expansion of EE with a doubling time ∼3h (a four-fold increase in 6h, Fig. 3F). This is consistent with previous studies suggesting that cell division is uniform across the early epiblast (*7, 8, 12*), and therefore cannot explain EE expansion on its own.

**Figure 3.**
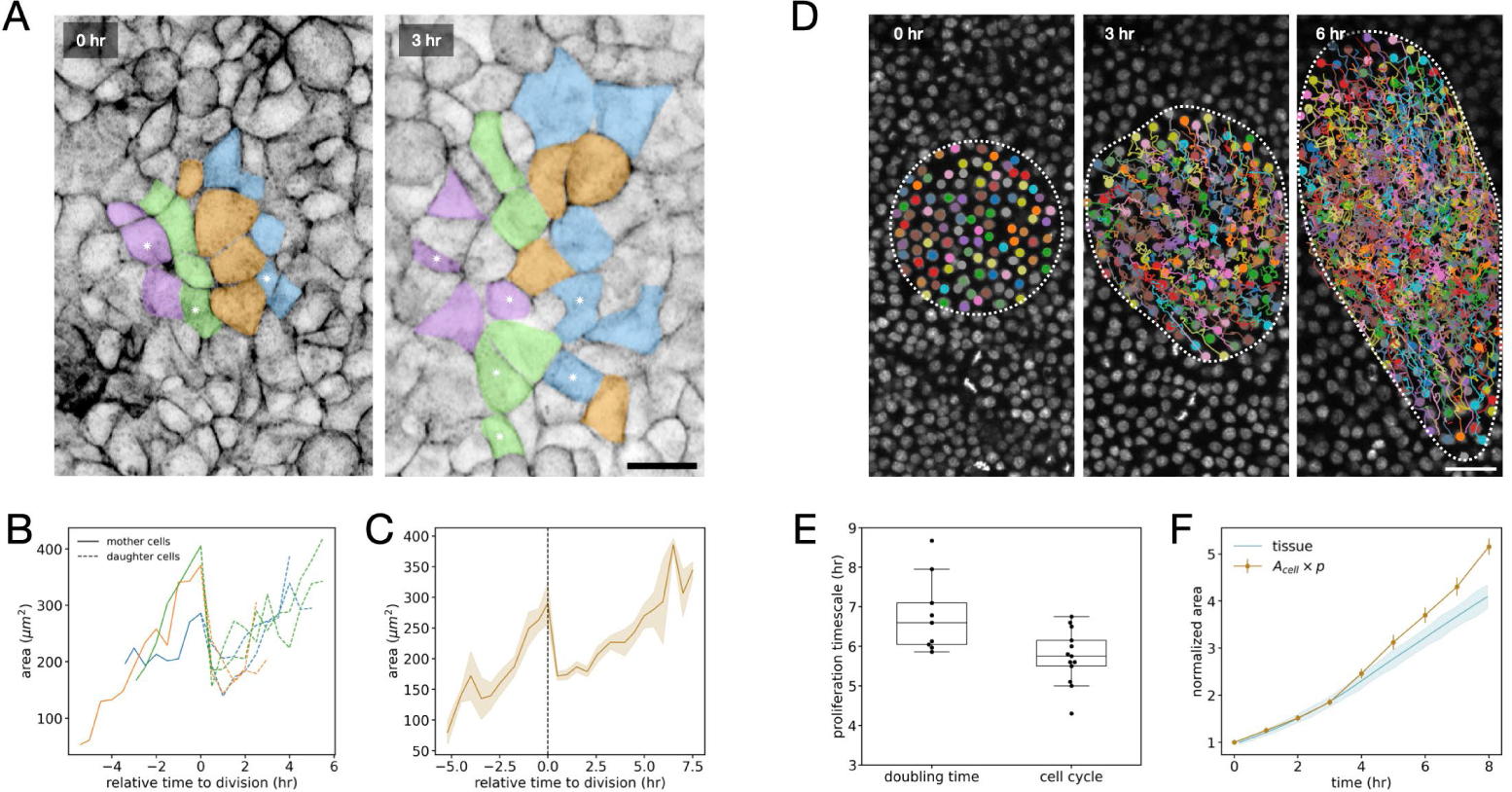
Cell division contribution to extra-embryonic expansion. (A) Manual tracking of a group of cells over 3 hours in a PIV-registered region. Dividing cells are identified by an asterisk. (B-C) Cell area quantification of dividing cells over time, aligned on the moment of their division for 3 individual mother cells (solid line) and their daughters (dashed line) in (B) and averaged, over n=10 mother cells and N=2 embryos in (C). (D) Automatic segmentation of nuclei and tracking in a PIV-registered region of interest. (E) Proliferation timescale, estimated by computing the cell doubling time in n=6 regions of interest from 4 embryos (N=6313 cells total) and by measuring the time between two consecutive divisions of n=13 tracked cells from N=2 embryos. (F) Evolution of the product of proliferation (measured with nuclei) with the cell area (measured with cell contours). Shaded regions and error bars correspond to the standard error of the mean. Scale bars: 20 µm (A), 40 µm (D).

### Mechanics of extra-embryonic tissue expansion

Since epiblast expansion has been shown to be driven by the migration of the epiblast border (*4*), we next asked whether cell area increase is caused by stretching. Whereas it is believed that the migration of the epiblast edge sets the entire disk under tension (*7, 8*), our data showing that embryonic and extra-embryonic regions deform differently suggested that the transmission of epiboly-induced tension may be restricted to the EE territory. To test this, we performed tissue-scale circular laser ablations (∼250 µm diameter) within the EP and EE territories (Fig. 4a). Since, for these large cutouts, the tissue relaxation time was comparable to the ablation time, we could not measure initial recoil velocities. Instead, we measured the total strain relaxation, which is a good indicator of relative tissue tension (*1*). We found that the area relaxation is much greater in the EE than in the EP, consistent with a greater state of tension (Fig. 4B). Importantly, since the same regions were imaged (and their deformation quantified) for 4 hours prior to ablation, we could compare their expansion before ablation to their relaxation after ablation (Fig. 4C,D). This revealed that the areal contraction upon stress relaxation is very close to areal expansion before ablation, arguing that epiboly-induced tension provokes a reversible stretching of the EE tissue, akin to the response of an elastic material.

**Figure 4.**
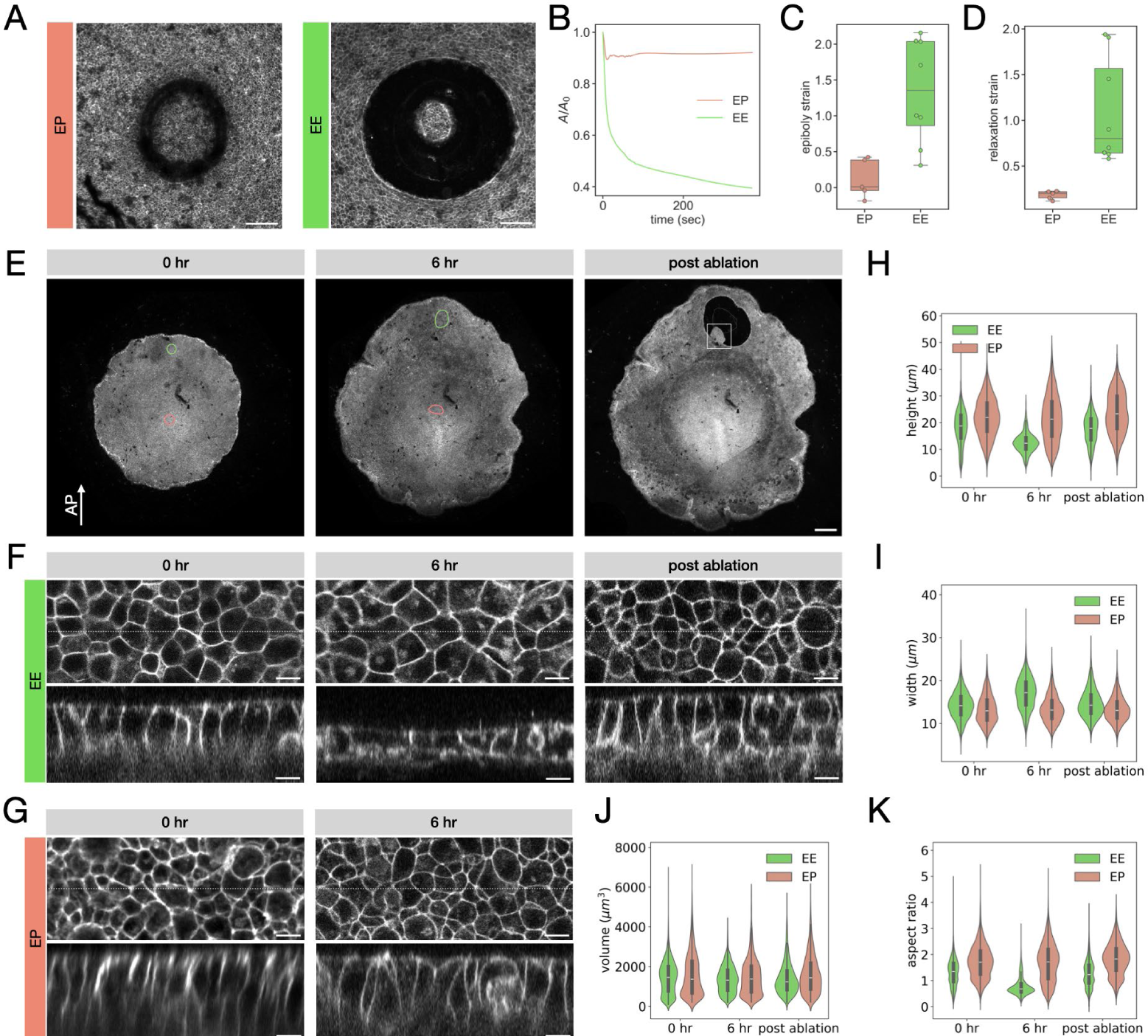
A reversible cell stretching accounts for the cell area increase. (A) Circular laser ablations in the embryo proper (left panel) and the extra-embryonic (right panel) regions, imaged 5 minutes after the laser cut. (B) Evolution of the ablated tissue area strain for the two regions displayed in (A). (C, D) Quantification of epiboly-induced strain, 5 hours before ablation (C) and relaxation strain, 5 minutes after ablation over n=13 regions from 5 embryos (D). (E) Time series of a transgenic memGFP embryo, in which two regions of interest are tracked by PIV in the embryonic (red) and the extra-embryonic (green) territories, before ablation and after ablation. (F, G) 2-photon imaging in regions of interest defined in (E) before the onset of epiboly (left panel), after 6 hours (central panel), and after laser-induced relaxation in the extra-embryonic region (right panel). Bottom row: XZ ortho-slices of the three-dimensional image at the location indicated by a dashed line in the top row. (H-K) Quantification of cell height (H), width (I), volume (J) and aspect ratio (K) over n=19,232 cells from N=3 embryos. Scale bars: 100 µm (A), 500 µm (E) and 10 µm (F, G).

To further characterize the response of the tissue, we sought to identify its cellular basis. We performed high-resolution 2-photon imaging to quantify the three-dimensional shape of cells before epiboly, after the onset of epiboly, and following laser-cut-induced tissue relaxation (Fig. 4E, Movie S5). These 3D acquisitions revealed that the initially columnar epithelial cells of the EE undergo an epithelial thinning as epiboly takes place and revert to their columnar conformation upon stress relaxation (Fig. 4F). This epithelial thinning is specific to the EE cells and does not occur in the EP, where cells retain their columnar morphology (Fig. 4G). To quantify this thinning process, we segmented cells in these regions using a custom ‘Cellpose’ model trained on our 3D dataset (Fig. S4). Quantifications showed that cell volume is approximately constant throughout the epiboly-induced deformation and laser-induced relaxation (Fig. 4H). The apicobasal length and the aspect ratio of cells remained constant in the EP, while they both decreased in the EE as epiboly took place, eventually reverting to approximately their initial values upon ablation (Fig. 4I-K, Table 1). Altogether these data reveal that columnar cells of the EE territory are reversibly stretched by the epiboly-induced tension into a squamous epithelial state. In terms of tissue material properties, this reversible morphological transition can be viewed as an ‘effectively elastic’ response (it is not strictly an elastic response since the remodeling of a tissue involves not only a passive deformation of its constituent cells but also an active remodeling of their contacts). Importantly, it does not take place within the EP, consistent with the epiboly-induced tension being restricted to the EE.

**Table 1.** Quantification of 3D segmentation of cells in the EP and EE regions.

|  | 0 hr |  | 6 hr |  | After ablation |  |
| --- | --- | --- | --- | --- | --- | --- |
|  | EP | EE | EP | EE | EP | EE |
| height ($\mu\text{m}$) | 21.9 | 18.2 | 21.6 | 12.5 | 23.7 | 17.5 |
| aspect ratio | 1.72 | 1.31 | 1.69 | 0.77 | 1.82 | 1.26 |
| volume ($\mu\text{m}^3$) | 1587 | 1450 | 1472 | 1358 | 1600 | 1400 |

### A simple viscoelastic model captures the morphogenesis of both embryonic and extra-embryonic territories

Our results show that the tension induced by the migration of the blastoderm edge induces a reversible, effectively elastic response in the EE territory, while the EP territory is mostly unaffected. Concomitantly, we have previously shown that the EP undergoes fluid-like motion that is driven by tensile stresses generated at the interface between the EP and EE territories and shapes the primitive streak. To explain how these driving forces and responses combine to give rise to differential area expansion in two contiguous epithelial territories, we considered a simple 2D viscoelastic model in which the tissue responds differentially by reversibly stretching under isotropic tension and irreversibly flowing under anisotropic (shear) stresses, with source terms for active tensile stresses at the margin and the border moving outward with a uniform velocity. In this framework, differences in tension along the margin account for the rotational tissue flows associated with primitive streak formation (*1*), while the average margin tension shields the EP from the epiboly-induced tension, restricting the areal expansion it induces to the EE (Fig. 5A). Based on this description, it is expected that the shape changes of tissue regions (associated with viscous shear flow) should be permanent, while area changes (associated with elastic expansion or contraction) should be reversible. To test this, we photoconverted regions in different parts of the epiblast using a mEOS2 photoconvertible transgenic quail line. In this way, we could track the deformation of photoconverted regions after 6 hours of development and their relaxation upon detachment of the entire circumference of the epiblast, which is impossible to achieve using PIV (since relaxation is too fast to be captured by PIV). As predicted by the model, regions in the EP exhibited little areal expansion and relaxation, regions in the EE stretched and reverted more significantly, and regions of the EE close to the margin fell in between. On the other hand, the aspect ratio of these regions remained unchanged upon relaxation, demonstrating their irreversible shape change. Notably, area relaxation in the EP and margin after ablation slightly exceeded the initial stretch (Fig. 5B,C, Movie S6), consistent with a build-up of tension at the margin when gastrulation is initiated (*1*). Taken together, these results provide a simple, unified model for the different mechanical behaviors of the entire epiblast, and show that the differential modeling of the EP and EE territories can be explained by the non-uniform mechanical state of the epiblast, with no need for tissue-specific responses.

**Figure 5.**
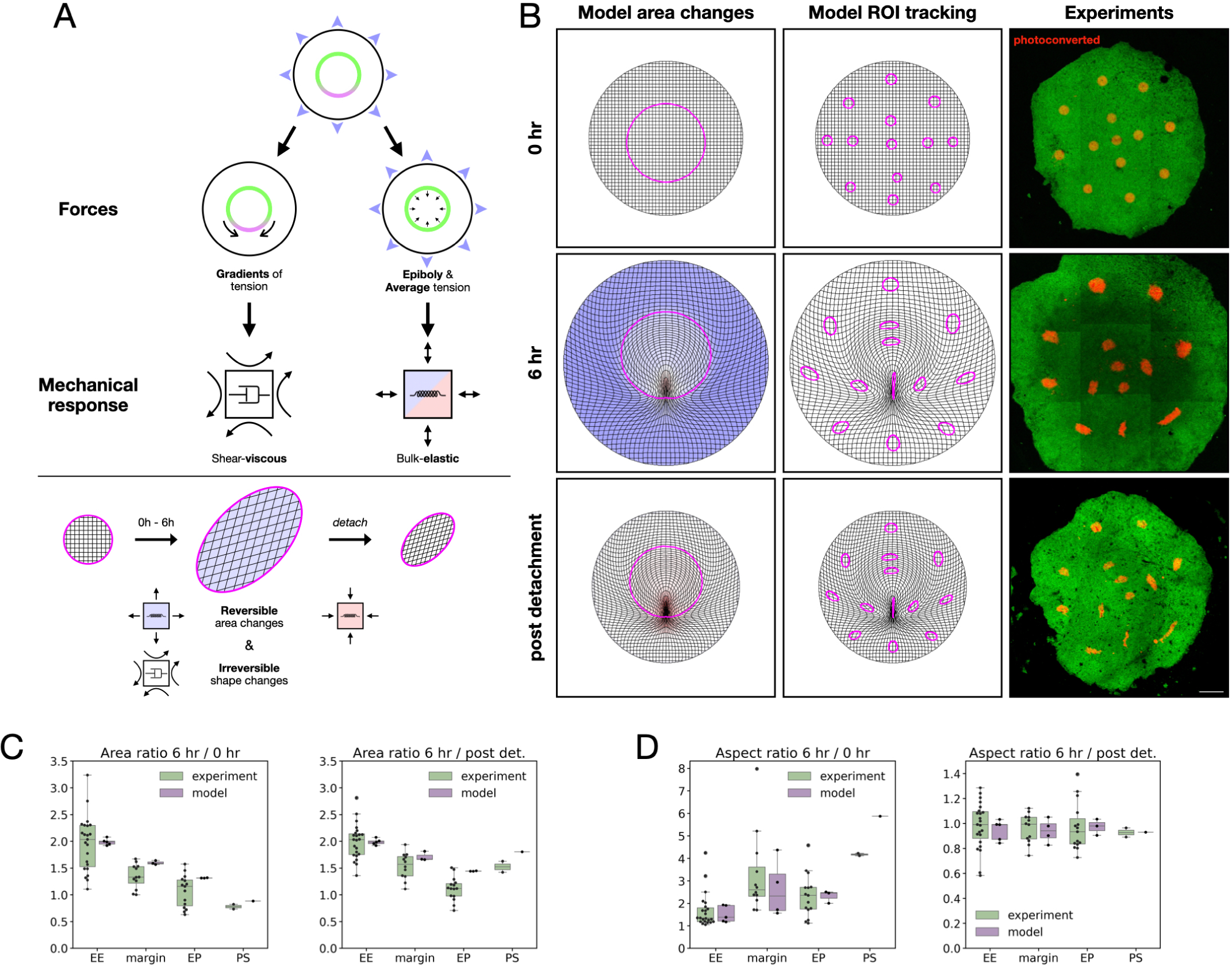
A viscoelastic mechanical model explains how epiboly and margin tension shape the entire embryo. (A) Schematic of the model. Upper panel: forces at work during the concomitant gastrulation and epiboly processes. Tension gradients (magenta) at the margin between the embryonic and extra-embryonic territories generate shear stresses across the epiblast, while the outward migration of the epiblast (blue arrows) produces a bulk extensile stress, that in the model is balanced by the average tension at the margin (green), which shields the EP from epiboly-induced stretching. The mechanical response of the tissue, which is uniform across the epiblast, is viscous for the shear and elastic for the bulk stresses. Lower panel: Deformation of the extra-embryonic territory induced by the combination of forces and mechanical response. In 6 hours of development, the EE both expands and flows. After detachment of the edge, the elastic expansion is reversed, while the viscous shape changes are not. (B) Comparison of model predicted deformation at 0h (top row), 6h (central row), and after detachment of the epiblast edge (bottom row) with experimental data. Left column: area change predicted by the model. Central column: deformations of regions of interest as predicted by the model. Right column: mEOS transgenic embryo with photoconverted regions (in red). (C, D) Quantification of growth and relaxation ratios for the area and aspect ratio of the regions of interest (ROIs) in the experiments and the model. (C-D) Area growth ratio (C) and aspect ratio change (D) between 6h and 0h, and relaxation ratio after detachment for n=52 ROIs (experiments), and n=12 ROIs (model). Scale bar: 500 µm.

## Discussion

In this study, we characterize epiblast expansion during avian epiboly. Our tissue-level analysis shows that the expansion is mostly limited to the EE territory. At the cellular level, apical area expansion is an essential contributor to EE growth. Cell areas eventually saturate and the increase in cell numbers must eventually become the predominant contributor, given the thousand-fold area increase necessary to cover the entire yolk. But cell division, which is uniform across the epiblast (*7, 8, 12*), cannot explain EE expansion on its own. Indeed, our quantifications show that proliferation is too slow to accommodate the rapid expansion of the epiblast, as imposed by the epiblast border migration; as proposed earlier (*8*), we conclude that the imbalance between proliferation and the migration of the edge cells leads to a state of tension.

It is common to analyze growth in a stationary regime, assuming a steady-state distribution of cell sizes. Then, growth can be viewed as intrinsic, and the direct consequence of proliferation (even where its rate is taken to vary in response to mechanical stresses). By contrast, our study of epiboly suggests a view where cells that stretch in response to tension serve as a reservoir of area, allowing the tissue to accommodate the expansion that is extrinsically imposed by the outward migration of its border. In this view, proliferation mostly serves to replenish a pool of smaller cells to sustain further expansion. This process may eventually settle into a stationary regime, especially if the expansion of the tissue vastly exceeds what the initial cells can support, but the distinction between intrinsically and extrinsically driven expansion remains.

We also clarify the process by which cells expand in response to tissue tension, showing that EE cells exhibit an effectively elastic response, changing reversibly their aspect ratio while keeping their volume constant. This result suggests that cell area expansion is solely a passive response to the extrinsic epiblast border migration, without contribution from a flattening mechanism intrinsic to the cells, at odds with previous suggestions (*13*).

Perhaps most intriguing is the difference between the EP and EE territories. The EP territory barely expands, and shows little relaxation upon laser ablation or whole epiblast border detachment compared to the EE, suggesting that the epiboly-induced tension does not propagate to the EP. This result runs against the common belief that the tension-induced epiboly is transmitted to both the EE and EP, although it is worth noting that this conclusion arose because only entire epiblasts were measured upon border detachment (*4, 7*). While it could be conceived that the differential expansion of EP and EE results from differences in their mechanical behavior, the contractile margin that acts as the engine of gastrulation movements (with net motion arising from a non-uniform tension) can also serve to shield the EP from the tissue tension in the surrounding EE. Indeed, we find that a viscoelastic model incorporating reversible area changes and irreversible shear flow, with a tensile ring and migrating boundary, can recapitulate the morphogenesis of EE and EP. The model accounts for the concomitant rotational tissue flows that shape the primitive streak and differential tissue expansion of EE vs. EP, with no need to modulate its parameters in space. It accurately predicts the shape and area changes upon detachment of the entire border, as well as a moderate relaxation of the EP (Fig. 5C), which has been reported experimentally in a previous study (*7*). Our model suggests that this relaxation may exceed the earlier expansion of the EP, due to the progressive tension buildup in the margin. Since tissue tension has been described to be required for proper development (*4, 7*), one could ask whether the embryonic defects reported in this study manifest a role of epiboly-induced tension in the regulation of contractility at the margin, which we have previously shown to be mechanosensitive (*14*).

Epiboly is an *in vivo* model of epithelial spreading during which tens of thousands of cells migrate collectively. As such, it can be compared with *in vitro* collective cell migration systems. Indeed, the expansion dynamics of systems with a similar geometry and size (disks of several mm) has been recently described (*15*). However, in these *in vitro* epithelial disks, growth is driven by proliferation, as opposed to the avian epiboly in which growth is driven by actively migrating edge cells. As a result, the timing between the two modes of growth is drastically different: patches of culture cells grow at the typical speed of 30 µm/hr while the embryo grows at 115 µm/hr, almost 4-fold faster. Given the critical nutritive function of the EE and the extensive yolk area it has to cover (10^9^ µm^2^) over 4 days, it is tempting to speculate that this fast growth mode by active migration has been selected during evolution.

## Acknowledgements

We thank Laure Bally-Cuif and Nicolas Dray for sharing their up-right microscope to image *in ovo* embryos. We gratefully acknowledge the UtechS Photonic BioImaging (Imagopole), C2RT, Institut Pasteur, supported by the French National Research Agency (France BioImaging; ANR-10–INBS–04; Investments for the Future), Laboratoire d’Excellence “Integrative Biology of Emerging Infectious Diseases” (Agence Nationale de la Recherche; ANR-10-LABX-62-IBEID, Investments for the Future). We especially thank Julien Fernandes for his help with 2-photon imaging. We also thank Marvin Albert, Gaëlle Letort and Léah Friedman for their help with image analysis.

This paper was typeset with the bioRxiv word template by @Chrelli: www.github.com/chrelli/bioRxiv-word-template

## Materials and Methods

### Animals

Animal husbandry for transgenic quails were carried out in accordance with the guidelines of the European Union 2010/63/UE, approved by the Institut Pasteur ethics committee authorization #dha210003, and under the GMO agreement #2432.

### Embryo imaging

Transgenic memGFP, H2B-GFP, mEOS quail embryos were collected at stage XI using a paper filter ring and cultured on a semi-solid nutritive medium of thin chicken albumen, agarose (0.2%), glucose, and NaCl, as described in (9). The embryos were then transferred to a bottom glass six-well plate (Mattek Inc.) with 2 mL (or 0.6 mL for high-resolution imaging) of the nutritive medium and imaged at 38°C with an inverted microscope Zeiss LSM 980 (except for ablation experiments and 2-photon imaging, see below) using 2.5X or 5X objectives (or a 20X for high-resolution imaging). The time interval between two consecutive frames was 6 minutes, except for nuclei tracking for which it was 3 or 4 minutes.

In the case of 2D cell contour imaging and nuclei tracking, Airyscan was used to obtain high-resolution images on a Zeiss LSM 980 microscope.

Regarding 3D cell contour imaging, embryos were imaged with a Zeiss LSM 7MP inverted 2-photon microscope using a Plan-Apochromat 40x objective at a wavelength of 850 nm owing to a Chameleon TiSaph laser. 2-photon imaging was also performed with a TrimScope Matrix II from Miltenyi Biotec using an Olympus 25X W/NA 1.05 objective at a wavelength of 850 nm owing to Spectra Physics Insight X3+ laser.

*In ovo* imaging was performed by transferring an egg full content to a 35-mm petri dish, closed with a Teflon membrane (FoilCover, Pecon). It was imaged with an upright microscope Zeiss LSM980 using a 2.5X objective. The time interval between two consecutive frames was 20 minutes.

### Laser ablation

Laser ablation was performed on an inverted microscope Zeiss LSM 900 using a UGA-42 firefly module coupled to a 355-nm pulsed laser (100% power) from Rapp Optoelectronic with a 10X objective. Images during ablation and the first minute following ablation were acquired every second. Images were subsequently acquired every 10 seconds.

### Embryo’s border segmentation

The embryo’s border was automatically detected by an active contour-based custom pipeline. In brief, the segmentation was initialized by manually defining the smallest circle around the embryo. The contour was then advected using the active contour method (skimage.segmentation.active_contour) provided by the scikit-image Python package (*16*). The advection parameters were manually defined for each dataset. The result of segmentation was then enlarged by a factor of 5% to initialize the segmentation for the following frame.

### Embryo’s border analysis

The embryo’s border cartesian coordinates (*x, y*) were transformed into a polar coordinates system (r, θ) with respect to the initial center of mass. To measure the border velocity, the time derivative of the distance to the center r was calculated at each angle using a second-order accurate central differences method.

### Quantitative analysis of tissue flows

Tissue flows were inferred from experimental movies using a previously reported custom particle image velocimetry (PIV) software, building on the ImageJ API for image preprocessing (*1*). This software provided the basis of our tissue quantification, in the form of frame-to-frame displacement grid data, the decomposition of observed flows into rotational and divergent components, and an automated snaking algorithm for the identification of the embryo proper (EP) contour based on contraction maps, and the embryonic edge based on intensity thresholding. For the decomposition of flows into radial and ortho-radial components, we defined radial polar coordinates with respect to the instantaneous barycenter of the EP contour. Radial and ortho-radial strain were computed by first computing the strain tensor in Cartesian coordinates using a centered finite difference on the PIV grid, and then transforming the components of the tensor to the instantaneously defined radial coordinates. EP and extra-embryonic areas were calculated using polygonal contours that were fit at intermediate stages and propagated through time using PIV. For quantitative comparison of strain and expansion rates across embryos, grid points were first sorted into 10 bins derived from a piecewise linear radial coordinate *u* such that *u* = 0 at the EP center, *u* = 1 on the EP contour and *u* = 2 on the edge. The data was then averaged in each bin before computing statistics across embryos

### 2D constitutive model of the embryo

We extend the fluid mechanical model of Saadaoui et al., 2020 by adding a non-linear bulk elastic response to the shear viscous behavior of the epiblast. Mathematically, the embryo is described as a homogeneous 2D material with constitutive relation

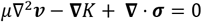

where *v* is the tissue velocity, *μ* is the shear viscosity, *σ* a localized stress on and tangential to the EP/EE margin, and *K* an elastic pressure that is a function of the pointwise net compression or stretch relative to the initial configuration. Denoting by *r* an effective basal length of epiblast cells that is calculated from accumulated area changes, we derive the elastic pressure *K*(*r*) = − *dv*/*dr* from a potential

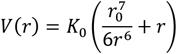

which models phenomenologically a strong resistance to compression, and a plateau in resistance to stretch. The stress *σ* derives from a line tension *T* = *T*_*base*_ + *T*_*dyn*_, where *T*_*base*_ is uniform along the margin, and *T*_*dyn*_ is larger in the posterior region. The boundary condition for the system is a prescribed radial velocity that of the embryonic edge based on experimental observations. Ablation of the edge is modelled by replacing this with a no-stress boundary condition for *t* ≥ *t*_*ablate*_ = 6h. The system is simulated numerically using the finite element package FEniCSx (*17, 18*), and the solution is post-processed using PIV for ROI tracking and quantification. Additional details may be found in the Supplementary Material.

### Regions of interest registration

Specific regions of interest were followed over time by running PIV on the movie. The region of interest was manually defined at a given time and the contour was then advected at all times of the movie using the flow field. The center of mass of the advected contour at each frame was calculated and used to register the movie.

### Cellular contour segmentation and quantification

memGFP contours were automatically segmented using a custom Cellpose model (*10*) trained on our dataset. We trained our custom model starting from the cyto2 Cellpose model, {size dataset} {parameters of the training}.

The segmentation output was loaded into the ImageJ ROI manager using a script inspired by the Cellpose imagej_roi_converter.py script. Using the ROI manager, the quality of the segmentation could be visually inspected. We estimated that an average of 95% of cells were correctly segmented. However, the quality of the segmentation was biased throughout the movie, as stretched cells become thinner and more transparent to ventral layers reducing the segmentation precision. To correct this bias, we only used for quantification a subset of frames (1 every hour) for which all cells had been manually proofread.

### Nuclei segmentation and quantification

Nuclei were segmented using the Stardist detector (*19*) directly integrated in Trackmate 7 (*20*). Tracking was then performed using the Advanced Kallman Tracker using mean intensity and mean area as weights. Using <u>Track Analyzer</u>, a tracking analysis Python package that we developed, we measured cell velocities and local area. In brief, the local area was computed by computing the Voronoi tessellation at each frame.

The number of cells in the region of interest was calculated by filtering out all cells outside the advected contour at each frame, using the Python shapely package (*21*). Proliferation was calculated by normalizing the number of cells by the initial number of each region of interest.

### Cellular 3D segmentation

memGFP 3D contours were automatically segmented using a custom Cellpose model (*10*) trained on our dataset. We trained our custom model starting from the TN1 Cellpose model on a dataset of 15 independent regions composed of roughly 40 cells from 3 different embryos. The training was performed on both XY images and XZ ortho-slices. The segmented 3D contours were analyzed by computing the convex hull for each cell using the scipy.spatial.ConvexHull method (*22*). For each cell, the convex hull was used to fit an ellipsoid and calculate the total volume.

### Proliferation quantification

The number of cells in the region of interest was calculated by filtering out all cells outside the advected contour at each frame, using the Python shapely package (*21*). Proliferation was calculated by normalizing the number of cells by the initial number of each region of interest. To compute the doubling time, we computed a linear regression on the log2 of the normalized cell number. Cell cycle was measured by manually tracking cells to ensure the quality of the tracking. The time between two successive divisions was measured.

To compute the product of proliferation by cell area, we multiplied the time evolution of the normalized number of nuclei averaged over all regions (from our H2B-GFP dataset) by the normalized cell area averaged over all regions (from our memGFP dataset).

### Ablation relaxation quantification

The area deformation of the ablated region prior to ablation was computed using our previously reported custom particle image velocimetry (PIV) software (*1*). We then calculated the pre-strain as (*A*_*f*_ − *A*_0_)/*A*_0_ with *A*_0_ and *A*_*f*_ being respectively the initial and final area of the region. In the case of the relaxation strain, we computed the evolution of the area of a patch of tissue within the ablated region, also using PIV. We then computed the strain as (*A*_*i*_ − *A*_∞_)/*A*_∞_ with *A*_*i*_ and *A*_∞_ being respectively the initial and final area of the patch of ablated tissue tracked by PIV.

### Photoconversion

Photoconversion of landmarks in the mEOS transgenic line was performed by illuminating 200 µm diameter disks with 80% laser power at 405 nm on a Zeiss LSM 900 with a 10X objective. A series of 30 iterations of roughly 1 second per region scanning was used to fully convert the green fluorescence into red fluorescence.

### Area quantification of photoconverted regions

To increase the contrast of photoconverted regions and minimize the effect of photobleaching, we calculated the difference between the red and green intensities. Then, we manually segmented the intensity difference.

## Supplementary Figures

**Figure S1.**
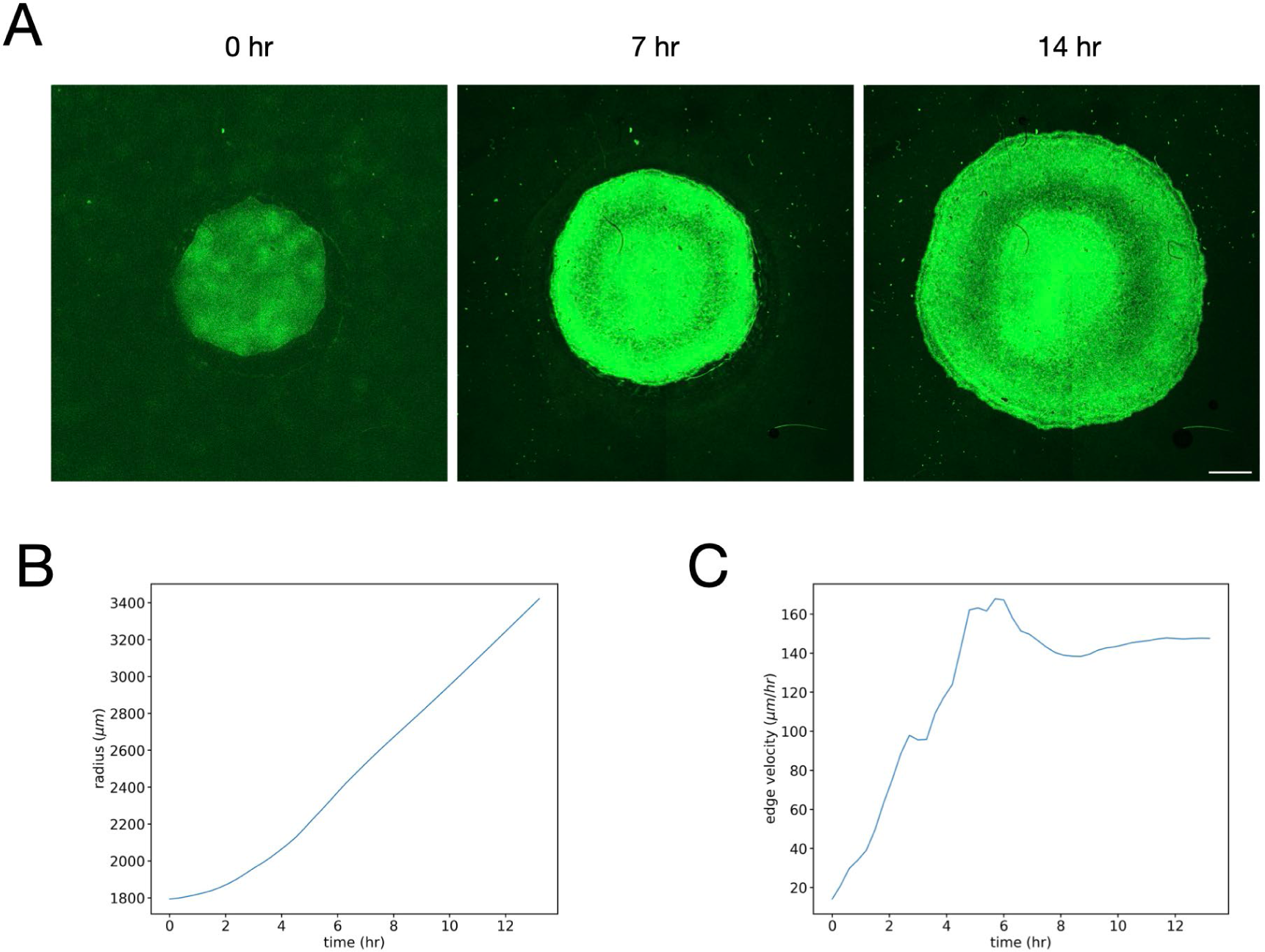
Quantification of epiboly in ovo. (A) Time series of a transgenic memGFP embryo live-imaged from stage XI over 14 hours. Scale bar: 1 mm. (B, C) Quantification of the embryo radius (B) and edge velocity (C) by automatic segmentation of the edge

**Figure S2.**
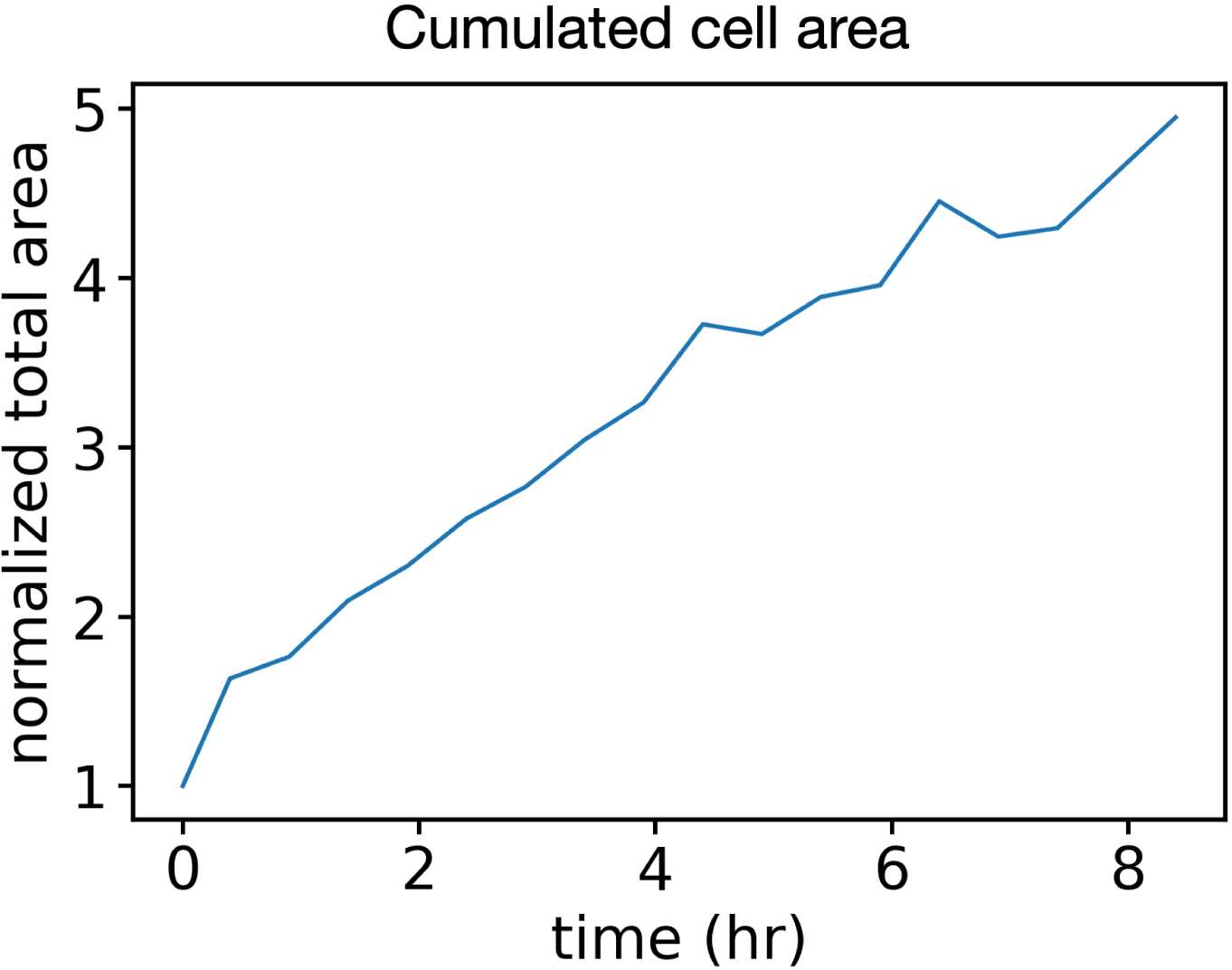
Evolution of the cumulated area of the mother cells and their daughter cells. The sum of the manually segmented cells’ area (Fig. 3 A-C) is computed and normalized by its initial value.

**Figure S3.**
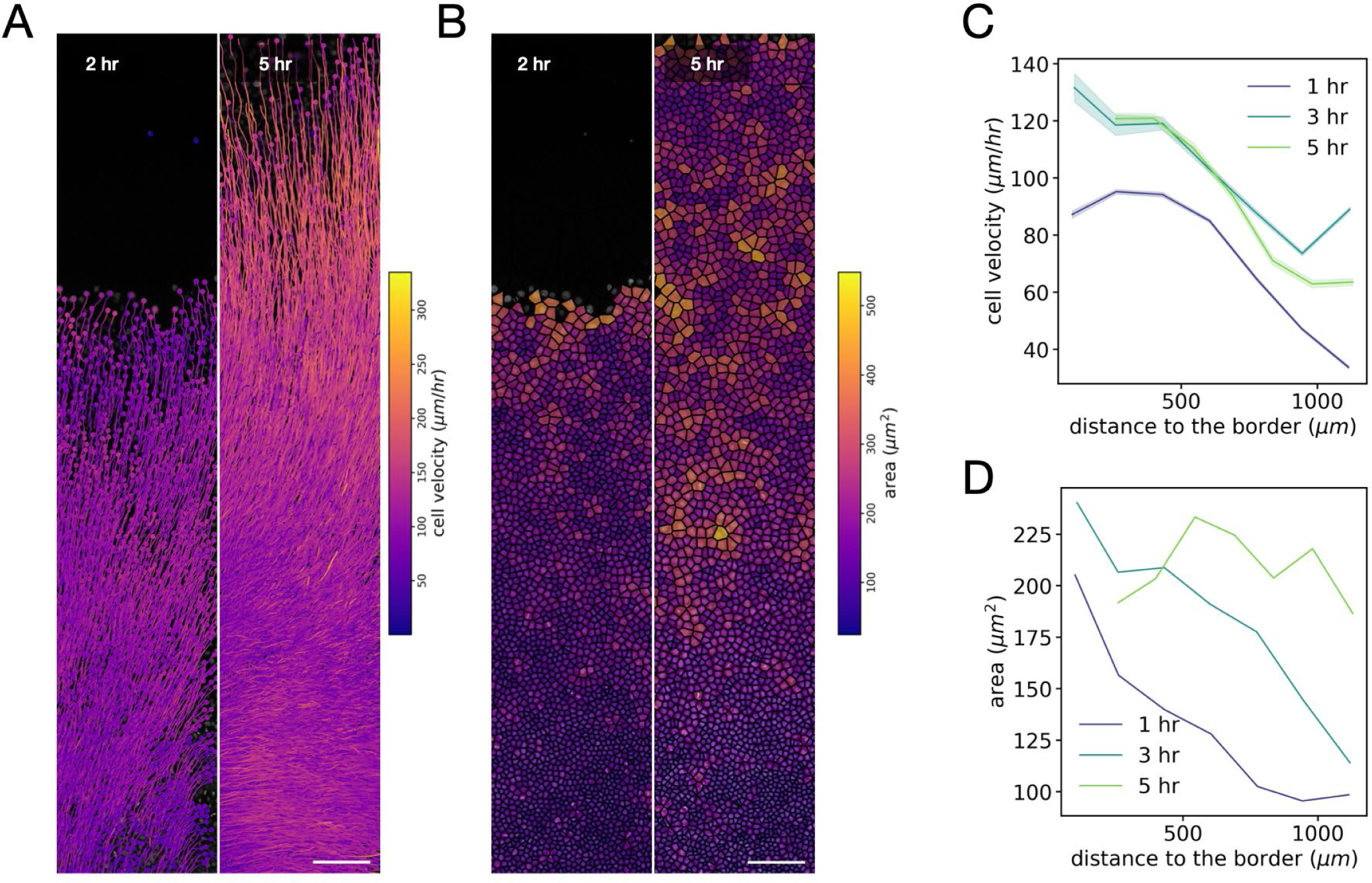
Quantification of nuclei tracking. (A-B) Time series of a strip of extra-embryonic tissue in a transgenic H2B-GFP embryo live-imaged from stage XI over 8 hours, overlaid with the velocity color-coded cell trajectories (A) and the area color-coded Voronoi tessellation (B). (C-D) Quantification of the cell velocity (C) and the Voronoi area (D). Cell velocity magnitude increases over time and shows a gradient with a maximum at the border with a value commensurate with the edge speed measured at low resolution. Similarly to our memGFP dataset, cell area initially shows a border-to-margin gradient, and then increases specifically in the central EE region.

**Figure S4.**
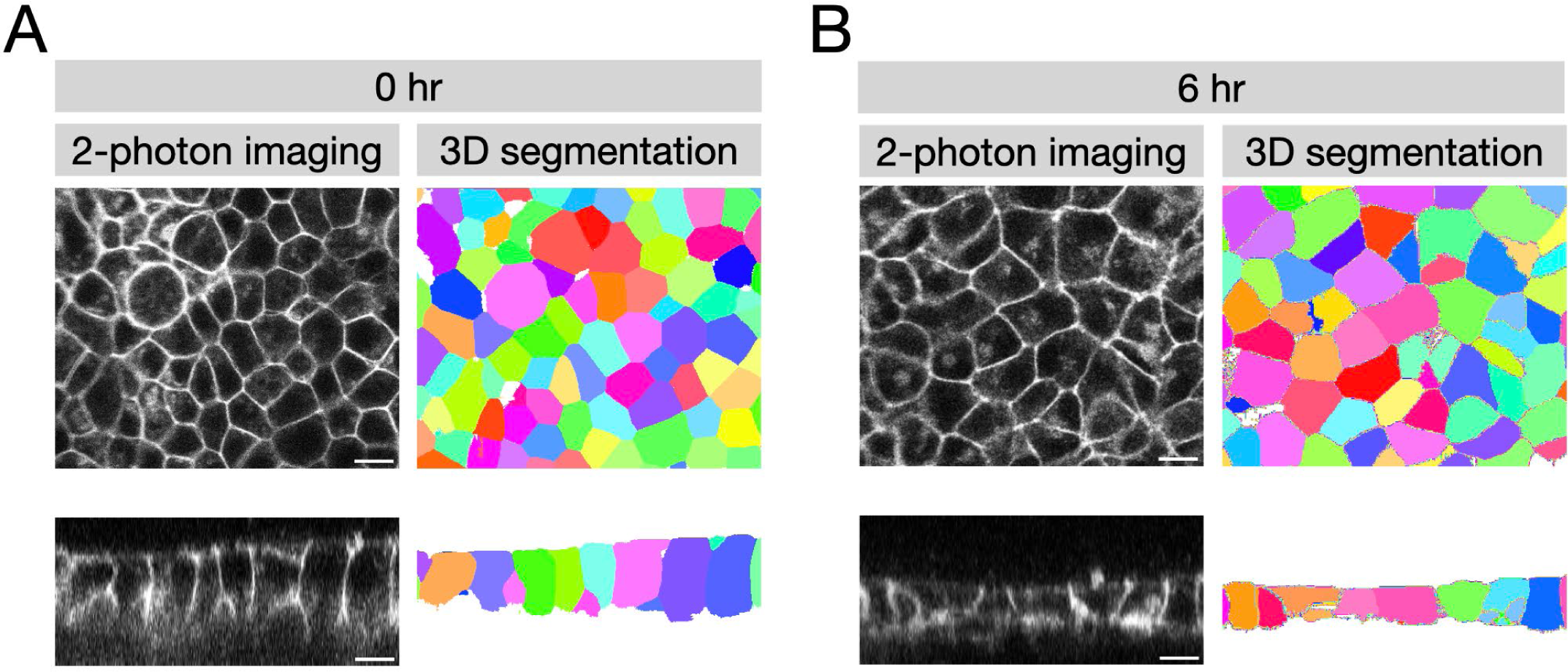
Quantification of three-dimensional cell volume and shape in the extra-embryonic. Cellpose segmentation of extra-embryonic cells at the onset of epiboly (A) and after 6 hours of epiboly (B). Left panels: original 2-photon images of XY slice (top row) and an ortho-slice (bottom row). Right panels: segmented cells color individually colored. Scale bars: 10 µm.

**Figure S5.**
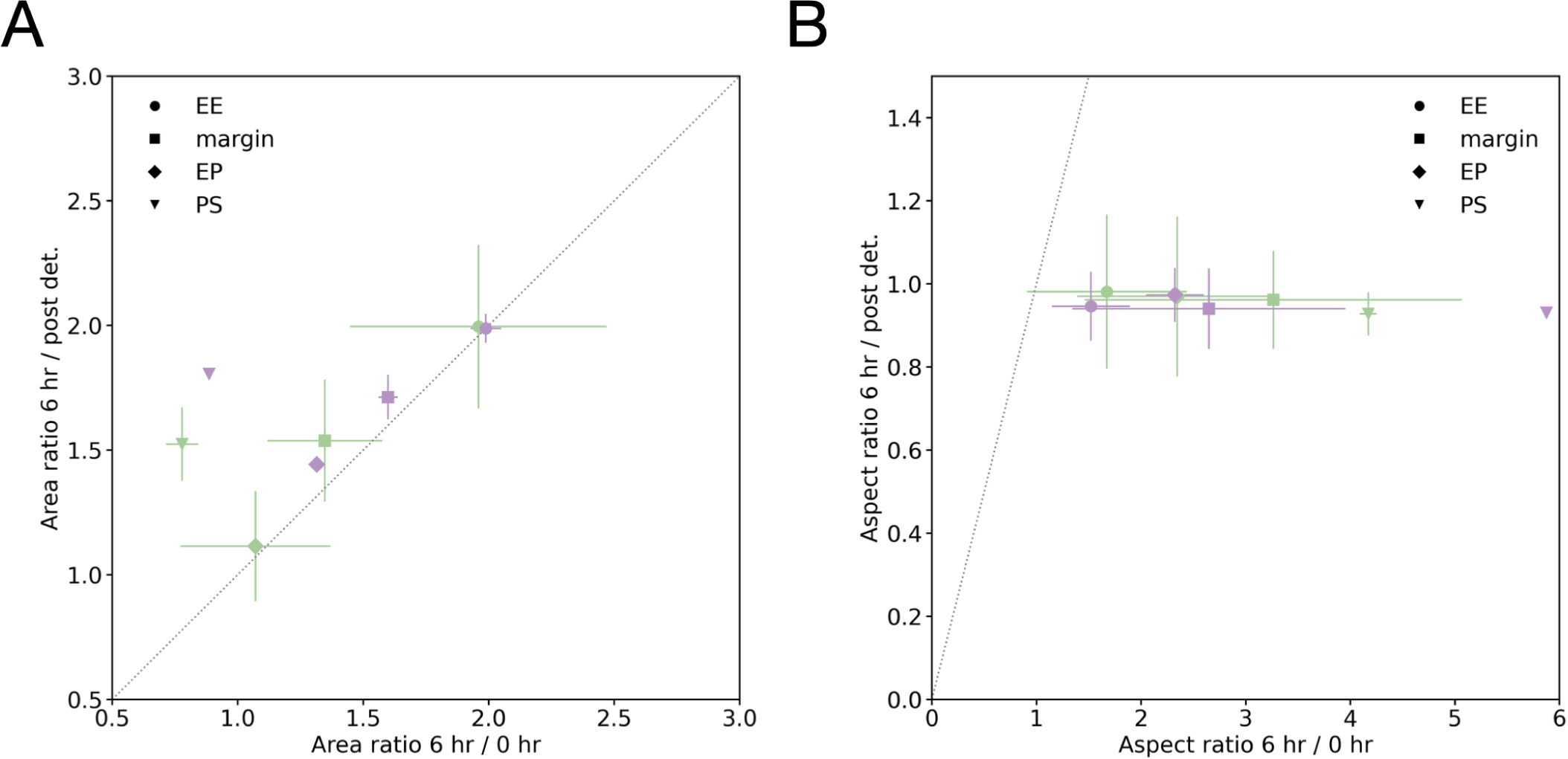
Comparison of expansion versus retraction of photoconverted regions before and after detachment of the embryonic edge. Left panel: Area ratio comparison. Green markers indicate mean and standard deviation of experimental data as in Fig. 5C, while purple markers indicate the same for sample ROIs calculated from the model. A dashed line indicates a 1:1 ratio. Points situated above the line indicate a more significant shrinkiage after detachment than expansion during the 6 hours before. Right panel: Analogous comparison for the aspect ratio. Points below the dashed line indicate a less significant change in aspect ratio after detachment than before.

## Supplementary movies legends

**Movie S1. Quantitative description of area changes during early quail embryo development**.

Timelapse movie of a memGFP transgenic embryo, capturing the concomitant gastrulation and epiboly tissue flows (upper right panel) and extraction of the divergent component of the flow (upper left panel) and further decomposition into radial and orthoradial strain rates (lower panels).

**Movie S2. PIV registration of regions of interest in the extra-embryonic territory**

High-resolution timelapse movie of a strip of extra-embryonic tissue in which 3 regions are tracked (left panel) and registered (right panel) using PIV to follow cell deformation, as epiboly is taking place.

**Movie S3. Quantification of extra-embryonic cell area during epiboly**

High-resolution time lapse movie of extra-embryonic cells (left panel) with their apical area color-coded (right panel) to highlight their expansion during epiboly.

**Movie S4. Manual tracking of a group of extra-embryonic cells during epiboly**

High-resolution timelapse movie in which a group of cells and their progeny is manually tracked in a PIV-registered region.

**Movie S5. PIV-based tracking of regions of interest before ablation**

Time lapse movie showing 2 regions in the EP and the EE territories tracked by PIV before laser ablation.

**Movie S6. Tracking tissue deformation of photoconverted regions before and after edge detachment**

Time lapse movie of mEOS2 transgenic embryo in which different regions of the epiblast are photoconverted (from green to red) to track their deformation before and after epiblast edge detachment.

