## Supplementary material for "A tension-induced morphological transition shapes the avian extra-embryonic territory"

### Supplementary Material for "Epiboly-induced tension stretches the avian extra-embryonic territory"

#### 2D CONSTITUTIVE MODEL OF THE EMBRYO

##### A. Model description

We extend the fluid mechanical model of Ref. [1] by adding a non-linear bulk elastic response to the shear viscous behavior of the epiblast. Mathematically, the embryo is defined as an initially circular 2D domain of radius $R_b$. Within this domain, a circle of radius $R_c$ and centre displaced posteriorly by a distance $d$ is defined as the initial EP/EE margin. The flows are calculated assuming homogeneous material properties with constitutive relation

$$\mu\nabla^2\boldsymbol{v} - \boldsymbol{\nabla}K + \boldsymbol{\nabla}\cdot\boldsymbol{\sigma} = \boldsymbol{0} \tag{1}$$

where $\boldsymbol{v}$ is the tissue velocity field, $\mu$ is the shear viscosity, $\boldsymbol{\sigma}$ a localized stress on and tangential to the EP/EE margin, and $K$ an elastic pressure that is a function of the effective 'basal length' $r$ of a cell located at position $\boldsymbol{x}$ in the embryo. The boundary condition for the system is a prescribed radial velocity $v_{\text{edge}}$ at the embryonic edge. Ablation of the edge is modelled by replacing this with a no-stress boundary condition for $t \geq t_{\text{ablate}}$.

The effective basal length $r$ is calculated from the accumulated deformation as follows. We denote the location of a material point in the undeformed initial configuration of the embryo by $\boldsymbol{X}$, and of the same point at time $t$ by $\boldsymbol{x}(t)$. The deformation vector is defined as $\boldsymbol{U} = \boldsymbol{x} - \boldsymbol{X}$, and the deformation gradient tensor $\boldsymbol{F}$ by $d\boldsymbol{x} = \boldsymbol{F}(\boldsymbol{X}, t)d\boldsymbol{X}$. Classical finite strain theory shows that the pointwise area change is given by the determinant $J = |\boldsymbol{F}|$. A straightforward algebraic manipulation then shows that

$$J = \frac{1}{|\boldsymbol{I} - \boldsymbol{\nabla}\boldsymbol{U}|}, \tag{2}$$

where $\boldsymbol{I}$ is the identity tensor and the gradient $\boldsymbol{\nabla}$ is understood to be taken with respect to the instantaneous (current) deformed configuration $\boldsymbol{x}$. With this in hand, we define the effective basal length as

$$r = r_0 \times \sqrt{J - a(t)m(\boldsymbol{x})}, \tag{3}$$

where $r_0$ is the initial basal length, assumed to be spatially uniform, and the square root accounts for the conversion of an area to a length. The additional term under the square root accounts for a spatially and temporally bounded prescribed loss of material at the primitive streak. This corresponds to the fifth mode of prescribed area changes in the synthetic embryo model of Ref. [1], and is the only one that is retained here. The expressions for $a(t)$ and $m(t)$ are given in the parameters section § B.

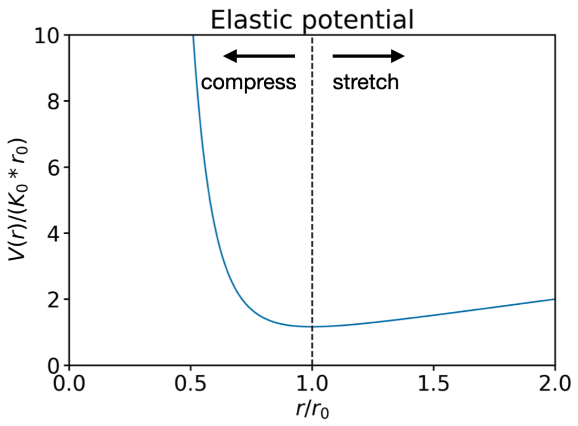


FIG. 1: Sketch of the elastic potential Eq. (4).

With this definition, we derive $K(r) = -dV/dr$ from a potential

$$V(r) = K_0\left(\frac{r_0^7}{6r^6} + r\right), \tag{4}$$

which models phenomenologically a strong resistance to compression, and a plateau in resistance to stretch. A sketch is shown in Fig. 1. Here $K_0$ is an overall scale for the elastic force and the equilibrium is at $r = r_0$. The form of this potential is in parts inspired by Ref. [2] who propose an elastic cell energy of the form

$$V(r) = K_0\left(\frac{2}{r^2} + r^4 - \frac{\alpha_l}{r} + \Lambda_\alpha r + \gamma_B r^2\right), \tag{5}$$

where the first two terms represent the nucleus resisting a too columnar or squamous shape, $\alpha_l$ is cell-cell adhesion, $\Lambda_\alpha$ is apical constriction (proportional to the cell perimeter), and $\gamma_B$ is basal adhesion to a substrate (proportional to basal area). The limiting behaviour both for small and large $r$ is governed by the nucleus terms. Removing the constraint on becoming squamous (which can be imagined to be relieved by cell divisions), and the effect of basal adhesion (which is absent in the epiblast, as a suspended epithelium), recovers the same $V \sim r$ behaviour for laterally stretched cells. For compressed cells,

| Parameter | Label | Value | Units |
|---|---|---|---|
| Initial EE radius | $R_b$ | 1.81 | mm |
| Initial EP radius | $R_c$ | 0.92 | mm |
| Initial EP posterior offset | $d$ | 0.106 | mm |
| Margin width | $w$ | 0.12 | mm |
| Epiboly velocity | $v_{\mathrm{ep}}$ | 0.15 | $\mathrm{mm\,h^{-1}}$ |
| Epiboly growth time scale | $t_{\mathrm{ep}}$ | 2.0 | h |
| Ablation time | $t_{\mathrm{ablate}}$ | 6.0 | h |
| Shear viscosity | $\mu$ | 1 | indet. |
| Equilibrium basal length | $r_0$ | 0.55 | $-^*$ |
| Bulk elasticity scale | $K_0$ | 10 | $\mu\,\mathrm{h^{-1}}$ |
| Baseline tension amplitude | $T_B$ | 3.0 | $\mu\,\mathrm{h^{-1}}$ |
| Maximal dyn. tension | $T_{\max}$ | 2.7 | $\mu\,\mathrm{h^{-1}}$ |
| Tension onset time | $t^+$ | 0.95 | h |
| Tension onset time scale | $\tau^+$ | 2.9 | h |
| Tension decay time scale | $\tau^-$ | 8.3 | h |
| Angular tension extent | $\Theta_T$ | 40° | – |
| PS location | $u_s$ | 0.945 | – |
| PS angular extent | $\Theta_s$ | 43° | – |
| PS width | $w_s$ | 0.13 | – |
| PS contraction | $a_0$ | 2.26 | – |
| PS contr. onset time | $t_s$ | 5.1 | h |
| PS contr. onset time scale | $\tau_s$ | 2.0 | h |

TABLE I: Numerical parameter values used in the simulation of the viscoelastic epiblast. Where relevant, the values correspond to the equivalent ones in Ref. [1]. The overall scale of the forces is an undetermined quantity, and relevant parameters have been defined relative to the shear viscosity. * The effective basal length in the calculation of the the elastic potential is thought of as defined relative to a fixed cell volume such that $r < 1$ indicates columnar, and $r > 1$ squamous shape.

we choose a steeper growth of the potential, which phenomenologically models our observation that the epiblast is nearly incompressible.

The stress $\boldsymbol{\sigma}$ is defined as

$$\boldsymbol{\sigma}(\boldsymbol{x},t) = T(s,t)\delta(\boldsymbol{x} - \boldsymbol{r}(\theta,t))\boldsymbol{t}\otimes\boldsymbol{t}, \tag{6}$$

where $\boldsymbol{r}$ is the contour of the EP/EE radius, and $\theta$ the polar angle measured from the posterior with respect to the instantaneous EP barycentre. The $\delta$-distribution is regularised by a Gaussian of width $w$ that is pointwise orthogonal to the tangent to the margin $\boldsymbol{t}$. The tension $T$ is composed of two contributions,

$$T(\theta,t) = T_{\mathrm{base}}(t) + T_{\mathrm{dyn}}(\theta,t), \tag{7}$$

where $T_{\mathrm{base}}$ is a uniform baseline tension, and $T_{\mathrm{dyn}}$ a dynamic tension profile that drives rotational gastrulation flows and is taken from the synthetic embryo model of [1]. Expressions for both are reported in the parameters section § B.

The system is simulated numerically using the finite element package FEniCSx [3, 4], and the solution is post-processed using custom PIV software for ROI tracking and quantification.

#### B. Model Parameters

The prescribed ingression of the primitive streak is modelled as in Ref. [1], which takes the form

$$m(u,\theta) = \exp\left(-\frac{(u-u_s)^2}{2w_s^2} - \frac{\theta^2}{2\Theta_s^2}\right), \tag{8}$$

$$\dot{a}(t) \propto \frac{1}{2}\left(1 + \tanh\frac{2\,(t-t_s)}{\tau_s}\right), \tag{9}$$

$$a(8\,\mathrm{h}) = a_0, \tag{10}$$

where $u$ is a piecewise linear radial coordinate from 0 at the instantaneous barycentre of the EP contour to 1 on the margin and 2 on the EE border.

The epiboly velocity is imposed as

$$v_{\mathrm{edge}}(t) = v_{\mathrm{ep}}\left(1 - e^{-t/t_{\mathrm{ep}}}\right), \quad 0 \le t \le t_{\mathrm{ablate}}. \tag{11}$$

The baseline tension evolves with time in a fashion that mirrors the ramp-up of epiboly over time, but is assumed to be maintained also after ablation,

$$T_{\mathrm{base}}(t) = T_B\left(1 - e^{-t/t_{\mathrm{ep}}}\right), \quad 0 \le t. \tag{12}$$

The dynamic tension is also defined as in Ref. [1],

$$T_{\mathrm{dyn}}(\theta,t) = \frac{T_{\max}}{2}\left(1 + \tanh\frac{2\,(t-t^+)}{\tau^+}\right)\exp\left(-\frac{t}{\tau^-}\right) \times \exp\left(-\frac{\theta^2}{2\Theta_T^2}\right), \quad 0 \le t. \tag{13}$$

All numerical values necessary to perform the simulation are listed in Table I. The most important choice for the application of the model to the elastic response after ablation is the ratio $\mu/K_0$, which sets the time scale of visco-elastic relaxation. Since the experimentally observed relaxation is fast, this ratio needs to be chosen small to account for this effect. We have found a value of $\mu/K_0 = 0.1\,\mathrm{h}$ sufficient to numerically capture relaxation to an approximately steady equilibrium state without interference from continuing gastrulation flows due to the continued presence of tension gradients in the margin.

[1] Mehdi Saadaoui, Didier Rocancourt, Julian Roussel, Francis Corson, and Jerome Gros. A tensile ring drives tissue flows to shape the gastrulating amniote embryo. *Science*, 367(6476):453–458, 2020.

[2] Edouard Hannezo, Jacques Prost, and Jean-Francois Joanny. Theory of epithelial sheet morphology in three dimensions. *Proceedings of the National Academy of Sciences*, 111(1):27–32, 2014.

[3] Martin S Alnæs, Anders Logg, Kristian B Ølgaard, Marie E Rognes, and Garth N Wells. Unified form language: A domain-specific language for weak formulations of partial differential equations. *ACM Transactions on Mathematical Software (TOMS)*, 40(2):1–37, 2014.

[4] Igor A Barrata, Joseph P Dean, Jørgen S Dokken, Michal Habera, Jack Hale, Chris Richardson, Marie E Rognes, Matthew W Scroggs, Nathan Sime, and Garth N Wells. Dolfinx: The next generation fenics problem solving environment. 2023.
